# A Deep Learning Approach to Estimate Collagenous Tissue Nonlinear Anisotropic Stress-Strain Responses from Microscopy Images

**DOI:** 10.1101/154161

**Authors:** Liang Liang, Minliang Liu, Wei Sun

## Abstract

Biological collagenous tissues comprised of networks of collagen fibers are suitable for a broad spectrum of medical applications owing to their attractive mechanical properties. In this study, we developed a noninvasive approach to estimate collagenous tissue elastic properties directly from microscopy images using Machine Learning (ML) techniques. Glutaraldehyde-treated bovine pericardium (GLBP) tissue, widely used in the fabrication of bioprosthetic heart valves and vascular patches, was chosen as a representative collagenous tissue. A Deep Learning model was designed and trained to process second harmonic generation (SHG) images of collagen networks in GLBP tissue samples, and directly predict the tissue elastic mechanical properties. The trained model is capable of identifying the overall tissue stiffness with a classification accuracy of 84%, and predicting the nonlinear anisotropic stress-strain curves with average regression errors of 0.021 and 0.031. Thus, this study demonstrates the feasibility and great potential of using the Deep Learning approach for fast and noninvasive assessment of collagenous tissue elastic properties from microstructural images.

## 1. INTRODUCTION

Biological collagenous tissues are comprised of networks of collagen fibers embedded in a ground substance [1, 2], which provide pliability and strength important for many normal physiological functions. The attractive biological and mechanical properties [3] also make collagenous tissues, mostly derived from animals as xenografts, suitable for a broad spectrum of medical applications such as bioprosthetic heart valve (BHV) [4, 5], cardiovascular grafting/patch [6, 7], tendon [8] and hernia [9] repair. However, due to the heterogeneity and inherent variability of biological tissues, the mechanical properties of collagenous tissues obtained at different locations even within the same individual (regardless whether animal or human) may differ, and may impact tissue-derived device function.

Many studies [10-16] have shown that the microstructure of soft tissues, particularly the collagen fiber network structure, is the key determinant of the tissue elastic properties at the macroscopic level. Advanced microscopy imaging techniques, such as second harmonic generation (SHG) imaging, has enabled noninvasive visualization of soft tissue collagen networks at the microstructural level. The elastic properties of collagenous tissues are traditionally obtained through destructive mechanical testing of harvested tissue samples (Figure 1). Ideally, the nonlinear anisotropic elastic properties of collagenous tissues could be directly estimated from noninvasive images (e.g. SHG images) of the tissue microstructure, such that xenografts could be carefully selected based on their mechanical properties and optimal, more predictable, tissue-derived device function could be ensured.

**Figure 1.**
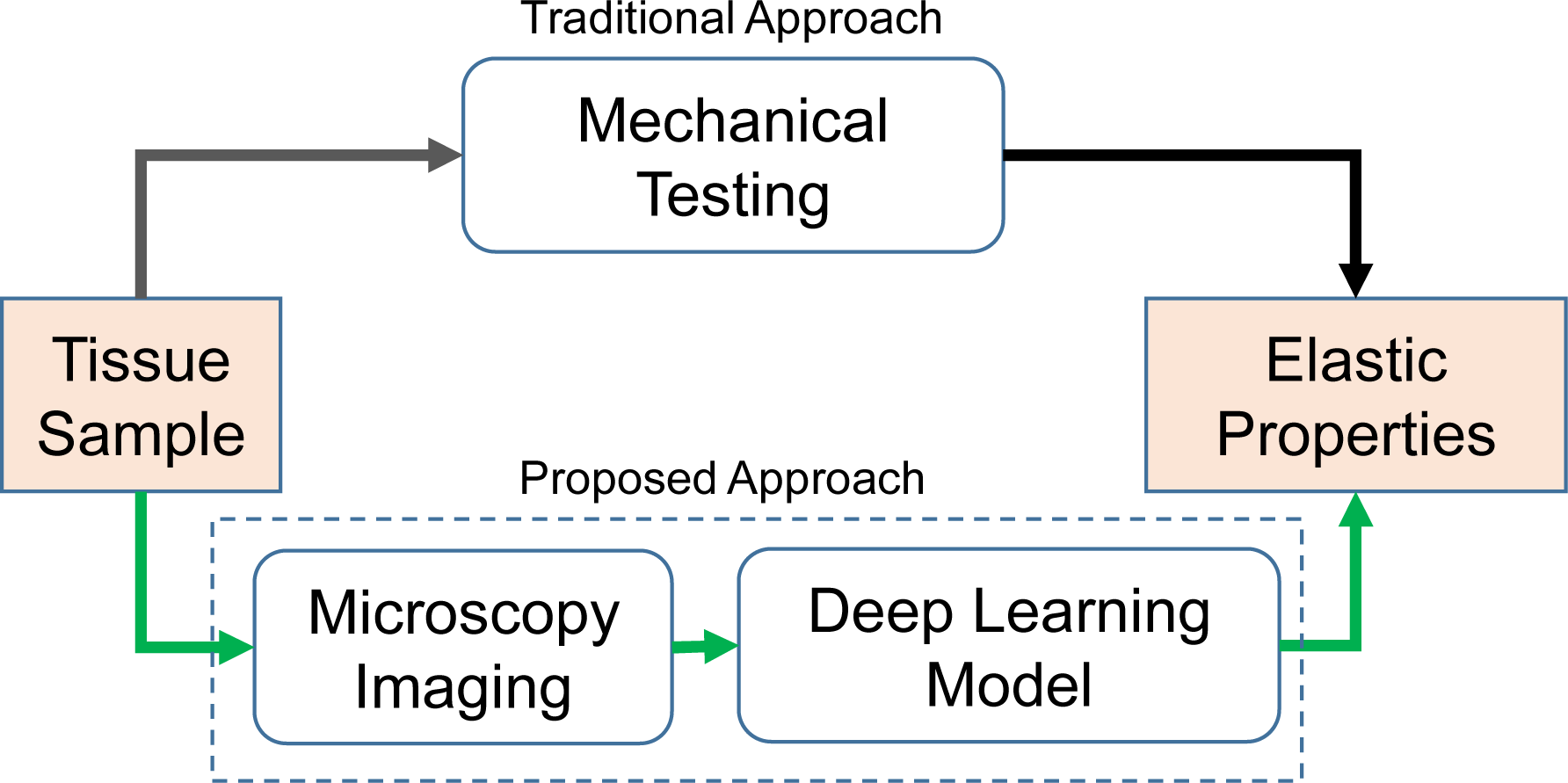
Two approaches to obtain the elastic properties of a tissue sample: 1) the traditional approach utilizing mechanical testing of a physical test sample and 2) noninvasive microscopy imaging coupled with a trained Deep Learning model.

Recently, Deep Learning [17], a branch of Machine Learning utilizing deep neural networks, has garnered enormous attention in the field of artificial intelligence. A special type of neural network, namely the convolutional neural network (CNN) [17-19], has become the state-of-the-art approach for computer vision and image analysis applications (e.g. face recognition), reaching, and even surpassing, human performance in some cases [20-23]. CNN provides an end-to-end solution from input image to output target value by automatically extracting image features, thus eliminating the need for hand-crafted image features.

In this study, we developed, to our best knowledge, the first Deep Learning approach to estimate the elastic properties of collagenous tissues from SHG images (Figure 1). Glutaraldehyde-treated bovine pericardium (GLBP) tissue, widely used in the fabrication of BHVs [5] and vascular patches, was chosen as a representative collagenous tissue. A multi-layer CNN was designed and trained on a dataset of SHG images and corresponding mechanical testing results (i.e., equi-biaxial stress-strain curves). The trained CNN can automatically extract features from input SHG images of GLBP tissues and predict the nonlinear anisotropic elastic properties (Figure 1).

## 2. METHODS

### 2.1 Tissue preparation and mechanical testing

The GLBP tissue samples used in this study were collected and mechanically tested through previous work by our group aimed at evaluating transcatheter heart valve biomaterials [24]. The tissue preparation and mechanical testing protocols are well documented in the published works [25-27]. Briefly, testing samples were cut into a 2×20 mm^2^ square, and four graphite markers delimiting a square approximately 2×2 mm^2^ in size were glued to the central region of each sample for optical strain measurements. Samples were then mounted in a trampoline fashion to a planar biaxial tester in aqueous 0.9% NaCl solution at 37 °C. A stress-controlled test protocol [25] was utilized to obtain the biaxial stress-strain response curves of each testing sample. In this study, 48 GLBP tissue samples were tested in total.

### 2.2 Tissue imaging

Upon completion of biaxial mechanical testing, the tissue samples were imaged using the SHG technique at the unloaded state. We utilized a Zeiss 710 NLO inverted confocal microscope (Carl Zeiss Microscopy, LLC, Thornwood, NY, USA), equipped with a mode-locked Ti:Sapphire Chameleon Ultra laser (Coherent Inc., Santa Clara, CA), a non-descanned detector (NDD), and a Plan-Apochromat 40x oil immersion objective. The laser was set to 800 nm and emission was filtered from 380–430 nm. Samples were kept hydrated with saline solution during imaging to prevent drying artifacts and covered with #1.5 coverslips. Samples were imaged inside the area delimited by the graphite markers, and 2D image slices were collected in the thickness direction from the smooth side of each sample. A 2D slice has 512×512 pixels to 1024×1024 pixels, and for each sample the number of slices was varied to cover the thickness. In total, we obtained 3D SHG images (size from 512×512×*N* to 1024×1024×*N*) of 48 tissue samples from different animal subjects, and the corresponding mechanical testing data. Representative SHG images of a GLBP sample are shown in Figure 2, with a total of 18 slices (*N*=18) through the thickness. It is evident from Figure 2 that the image patterns change very slowly through the GLBP tissue thickness.

**Figure 2.**
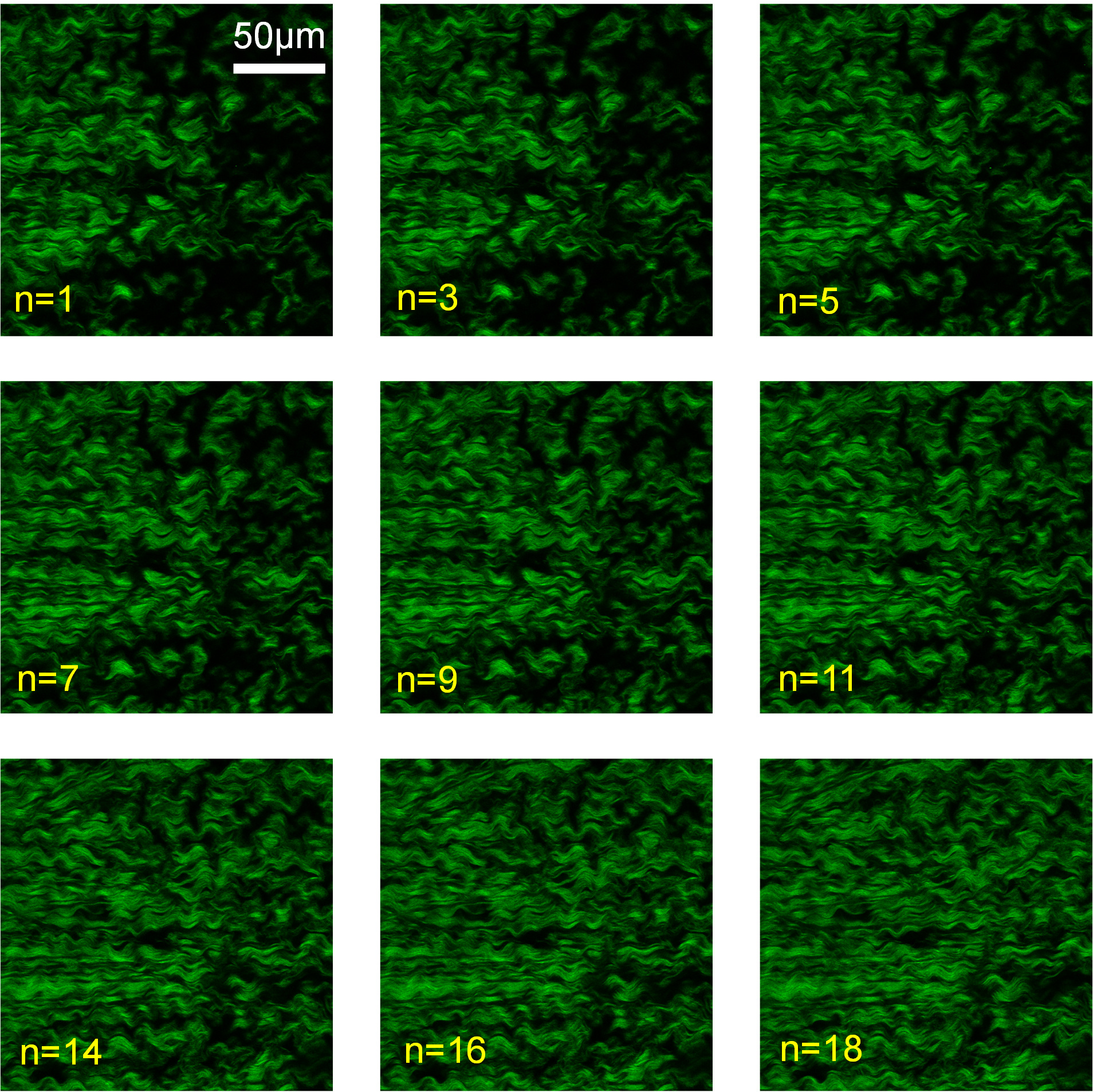
Representative SHG image slices of a tissue sample. n denotes the index of each slice.

### 2.3 PCA-based Parameterization of GLBP stress-strain curves

Two distinct stress-strain curves were obtained from the equi-biaxial mechanical testing (section 2.1) of each tissue sample (Figure 3a&b), due to the anisotropic mechanical behavior of the tissue: 1) strain E_11_ and stress S_11_ along the X_1_-direction, and 2) strain E22 and stress S22 along the X_2_-direction. Each stress-strain curve was uniformly sampled along the stress axis within the range of 10 to 630 *KPa.* The cutoff of *630KPa* was chosen because different ranges of external stresses were applied to the tissue samples and *630KPa* was the minimum peak stress value. For each tissue sample, the resampled strain values from the two curves were assembled as a vector of 126 numbers, *Y*. By using principle component analysis (PCA) [28, 29], the vector *Y* of a tissue sample can be decomposed as
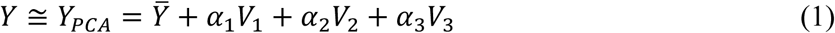

where 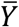 is the population mean, {*V_i_*} are the modes of variation, and {*α_i_*} are the coefficients. Here, {*α_i_*} can vary, while 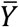 and {*V_i_*} are the same for all tissue samples. The first three modes of variation {*V*_1_, *V*_2_, *V*_3_} with {*α*_1_, *α*_2_, *α*_3_} can describe 99% of the total variation of the stress-strain curves, which means each stress-strain curve can be almost perfectly reconstructed by using Eq.(1) as shown in Figure 3b. Furthermore, the reconstruction error was measured by the mean absolute error (MAE), given by
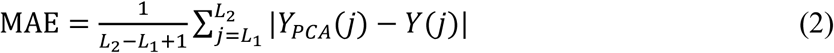

where *j* is the index of a component in a vector; and if *L*_1_ = 1, *L*_2_ = 63, MAE is the error of the reconstructed S_11_~E_11_ curve; and if *L*_1_ = 64, *L*_2_ = 126, MAE is the error of the reconstructed S_22_~E_22_ curve.

**Figure 3.**
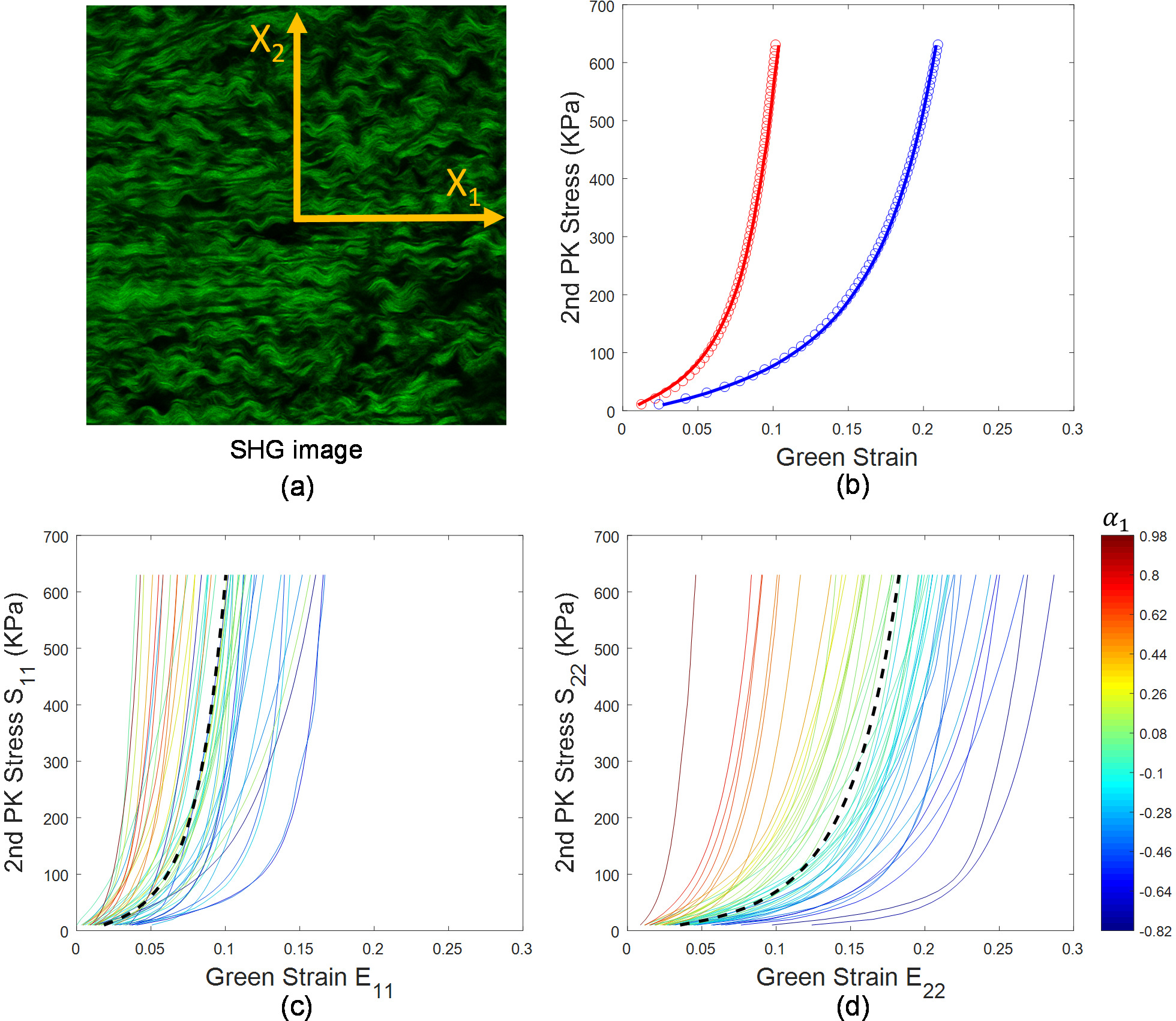
(a) The orientation definition of a tissue sample: X_1_ direction and X_2_ direction. (b) The open circles represent the stress-strain curves of a tissue sample from equi-biaxial mechanical testing experiments. The reconstructed stress-strain curves are shown by the red lines (S_11_~E_11_) and blue lines (S_22_~E_22_). (c)&(d) The stress-strain curves in the two directions of the 48 tissue samples color-coded by the corresponding *α*_1_. The dashed lines are the mean curves, 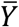.

As shown in Figure 3c&d, a material is softer, i.e. more compliant, than the mean material if *α*_1_ < 0, and stiffer than the mean material if *α*_1_ > 0. Thus the sign of *α*_1_ can be used to describe the overall tissue stiffness.

### 2.4 Deep learning model

As show in Figure 4, we designed a deep convolutional neural network (CNN) as the deep learning model, consisting of 6 blocks in a pipeline. The 1^st^ block takes an input image of size 256×256×N pixels. The 6^th^ block can be configured either as a classifier of the overall tissue stiffness (sign of *α*_1_), or a regressor to predict the PCA parameters {*α*_1_, *α*_2_, *α*_3_}, which can be used to reconstruct the stress-strain curves by Eq.(1). The CNN (Figure 4) learns the relationship between the tissue SHG images and elastic properties from the training dataset, and then can infer the elastic properties from a new tissue image.

**Figure 4.**
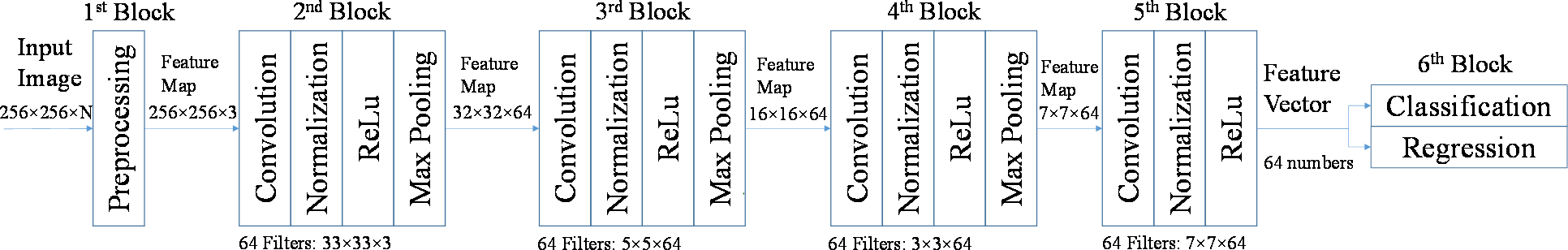
Architecture of the deep convolutional neural network used in this study.

Usually, convolutional neural networks (CNNs) consist of many layers that are sequentially connected, e.g., output from the first layer is the input to the second layer. A layer performs a specific operation, such as convolution, normalization, or max pooling, and it has parameters either prescribed or to be learned from data. For a detailed explanation of these layers, we refer the reader to the reference papers [17, 18, 30, 31]. The network structure should be designed for specific applications, e.g., choosing the types and sizes of layers and determining their combinations. For our application, the designed CNN consisting of 6 blocks in a pipeline, where each block has one or more layers. Given an input 3D image of 256×256×*N* pixels, the 1^st^ block with only one preprocessing layer, performs local contrast normalization and uniformly resamples the input 3D image into the first feature map of 256×256×3 pixels. The 1^st^ block does not have any trainable/free parameters. The 2^nd^ block contains a convolution layer with 64 filters (a.k.a. kernels) of 33×33×3 pixels, a batch-normalization layer, a ReLu (rectifier linear unit) layer, and a max pooling layer; and the output from the 2^nd^ block is a feature map of 32×32×64 pixels. The 3^rd^ to the 5^th^ blocks are very similar to the 2^nd^ block, which output feature maps of 16×16×64, 7×7×64, and 1×1×64 pixels respectively. All of the max-pooling layers use a 2×2 pooling window. The 1^st^ to 5^th^ blocks can be considered image-feature extractors which output a feature vector of 64 numbers. The 6^th^ block is used for classification with a softmax classifier, and regression with a linear model. The CNN was implemented by using MatConvnet [32], an open source MATLAB toolbox, and custom MATLAB functions; and it can process an input 3D image within 10 seconds on a PC with intel i7-4770 CPU and 32G RAM.

### 2.5 Learning of the deep convolutional neural network

The CNN (Figure 4) parameters were learned from the training data. To overcome the challenge of training the CNN with a small dataset [28] (i.e., 48 test samples, which is an acceptable sample size for material testing of biological tissues), the CNN was trained by combining: 1) unsupervised deep learning to determine the parameters in the 2^nd^ to 5^th^ blocks, 2) supervised learning to determine the parameters in the 6^th^ blocks, and 3) data augmentation to generate more training data.

#### 2.5.1 Unsupervised Deep Learning from the 2^nd^ to 5^th^ blocks

To determine the filter parameters of a convolution layer, generally we could use encoder-decoder based unsupervised learning strategies [33-36]. The input feature map to the convolution layer can be divided into small patches, where each patch has the same size as a filter (all filters in the same layer have the same size). Each patch can be converted to a vector, *X*, and the vectorized patches can be stacked together as the columns of a data matrix ***X***. The filters of the convolution layer can also be vectorized and stacked together as the columns of a filter matrix ***A***. Let *h*(*x*) denote the ReLu function: *h*(*x*) = *x* if *x* > 0, and *h*(*x*) = 0 if *x* < 0. The encoder performs convolution followed by ReLu to each patch, which outputs the code matrix *h*(***AX***) close to the optimal (unknown yet) code matrix ***Z***. Given the optimal code matrix ***Z***, the decoder tries to recover the input patches ***X*** by using a linear combination of the atoms/columns in a dictionary/matrix ***D***, i.e, using ***DZ*** to approximate ***X***. Then the goal is to find the optimal variables {***A*, *D*, *Z***} such that the encoding error and the decoding error are both minimized, which is to minimize the following objective function:
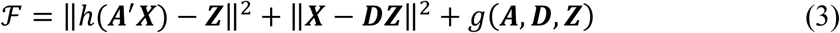

where *g*(***A*,*D*,*Z***) defines some constraints on the variables, and *A’* denotes the matrix-transpose of ***A***. The matrix norm **|**|. **|**| is the Frobenius norm. Obviously, by using different constraints, we can obtain different solutions of {***A***, ***D*, *Z***}. We proposed an algorithm with three steps to directly obtain a solution under the low rank constraint [37]:

*<u>Step-1</u>:* Perform low rank approximation (LRA) [37] on the patches ***X***, then a vectorized patch *X* can be approximated by
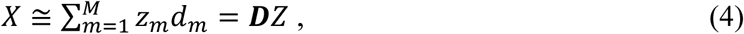

where ***D*** = [*d*_1_, …, *d_M_*], and the vector *d_m_* has the same size as *X*, and 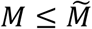 which is the number of pixels in the patch *X*. *d_m_* is the product of the *m*^th^ largest singular value, λ*_m_*, and the corresponding left-singular vector obtained by LRA. ***D*** is the same for every single patch *X*. Also obtained by LRA, the code vector, *Z* = [*z*_1_,… *z_M_*]’ is a column vector of scalars, which is different for different patches. The percentage error of approximation for the patches ***X*** is given by
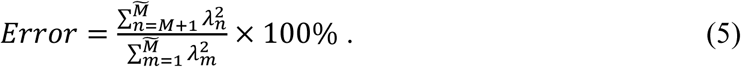

If 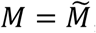, then the error is zero. By controlling the number of retrained singular values and singular vectors, i.e., *M*, the approximation error and the computation cost (proportional to *M*) can be controlled. In this study, *M* is fixed to 32, and the error is less than 30%. The low rank approximation essentially obtains ***D*** and ***Z*** that minimize ║***X– DZ***║^2^ under the low rank constraint. Since the singular vectors in ***D*** are orthogonal to each other, the code vector *Z* can be simply approximated by ***D****′X,* i.e., *Z* ≅ ***D****′X*, which is obtained by multiplying ***D***′ to both sides of Eq.(4). After this step, the code matrix ***Z*** and dictionary ***D*** are determined.

*<u>Step-2</u>:* Define the filter matrix ***A*** by using the learned dictionary ***D***, given by
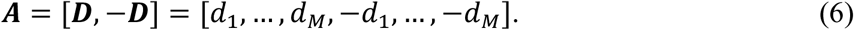

Also, we define a new code vector 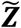 as
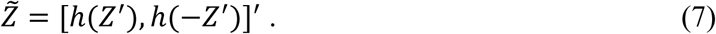

Then the objective function Eq.(3) is equivalent to
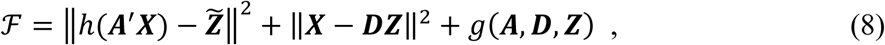

where 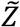 is the stack of new code vectors, corresponding to ***Z***, and 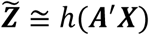 because *Z* ≅ ***D***’*X*. Then, a vectorized patch *X* can be encoded as a vector 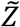 by the encoder *h*(***A***’*X*). For example, *X* = 2*d*_1_ − *3d*_2_, then the code vector is [2,0] if ***A*** = [*d*_1_, *d*_2_], and the code vector is [2,0,0, 3] if ***A*** = [*d*_1_, *d*_2_−*d*_1_,−*d*_2_], which clearly shows that the longer code vector preserves more information of *X*. The rationale of Eq.(6)&(7) is that the ReLu layer rejects any negative signal (i.e. code) output from the convolution layer, and therefore, nearly half of the signals will be lost in each block, harming the performance of the CNN. After this step, the filters of the convolution layer are determined.

*<u>Step-3</u>:* Perform feature map normalization. The output from the ReLu layer is a feature map serving as the input to the next layer. The size of the feature map is *K*_1_ × *K*_2_ × *K*_3_ (i.e. height × width × channel). The values of the feature map at one spatial location can be assembled to a code vector *Z* of length *K*_3_. By assembling all of the code vectors from the training dataset, a data matrix is obtained, and each row of this matrix is normalized by subtracting the mean and dividing by the standard deviation. The rows of the code matrix ***Z*** from a single input image will also be normalized in the same way by using the same values of mean and standard deviation. This normalization is essentially equivalent to batch-normalization [30] which has been shown to improve CNN accuracy. After this step, the parameters (i.e. mean and standard deviation values) of the normalization layer are determined.

#### 2.5.2 Supervised learning in the 6^th^ block

The 6^th^ block can be configured either as a classifier or regressor. In the classification configuration, a softmax function is used to predict class membership based on the feature vector from the 5^th^ block. Since it is a binary (soft vs. stiff) classification task, the softmax function reduces to a logistic function, given by
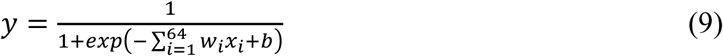

where {*w*_1_,…,*w*_64_, *b*} are the unknown scalar parameters and [*x*_1_,…,*x*_64_] is the feature vector from the 5^th^ block. Usually, a discrimination threshold (e.g. 0.5) is specified for the binary classification. If *y* is greater than or equal to the threshold, then the input is classified as stiff; and if *y* is smaller than the threshold, then it is classified as soft. With the labeled training data (i.e., image data with known mechanical properties), the 65 parameters in Eq.(9) can be determined through supervised learning using the cross-entropy loss function and the conjugate gradient optimization algorithm.

In the regression configuration, a multiple output linear regressor predicts the values of {*α*_1_, *α*_2_, *α*_3_} in Eq.(1) based on the feature vector from the 5^th^ block, which is given by
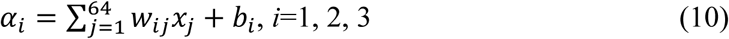

where {*w_i j_*,*b_i_*, *i* = 1,2,3, *j* = 1,…,64} are the unknown scalar parameters. With the labeled training data, the 195 parameters of this regressor can be learned by using the least squares regression algorithm. Once the parameters {*α*_1,_ *α*_2_, *α*_3_} are predicted by the regressor, the stress-strain curves can be reconstructed by using Eq.(1).

#### 2.5.3 Data augmentation

Data augmentation methods are extensively used in Deep Learning applications [18, 38-40] to generate more training data. In this study, two data augmentation methods were used: image splitting and flipping (Figure 5). A 3D image of *N* slices can be split into patches using a sliding window with a stride of 128, and the size of each patch is 256×256×*N*. As a result of image splitting, 1678 patches were generated. Furthermore, each patch was flipped along the horizontal direction and/or vertical direction, which produced 6712 patches. The elastic properties corresponding to image patches from the same GLBP tissue sample, were assumed to be identical.

**Figure.5.**
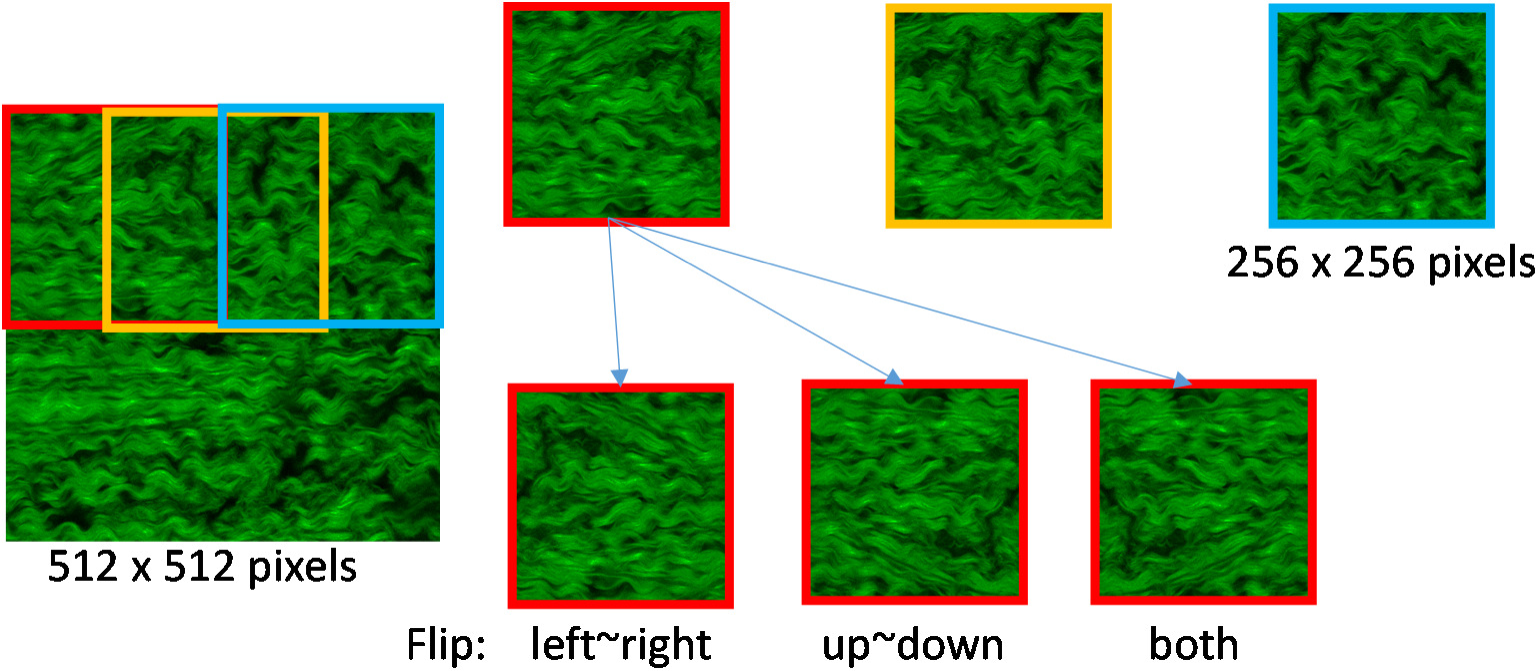
An example of data augmentation to generate image patches.

### 2.6 A comparative study of network structures

Given the relatively large size of the CNN compared to the dataset, the natural question arises whether reducing the number of layers or filters will significantly impact the performance. Given the huge design space, it would be impractical to evaluate all possible simplifications of the CNN structure. In this study, we chose to investigate two simplified CNNs for comparison, named CNN-s1 and CNN-s2 respectively. *<u>CNN-s1</u>*: in *Step-2* of unsupervised learning in section 4.5.1, the filter matrix ***A*** was simplified as ***A*** = ***D***, which reduces the number of filters. <u>*CNN-s2*:</u> the ReLu and normalization layers were removed, and the filter matrix ***A*** was simplified as ***A* = *D***. The structure of CNN-s2_is similar to that in [34].

## 3. RESULTS

### 3.1 Unsupervised deep learning

The learned filters of the CNN are visualized in Figure 6. The filters in the 2^nd^ block (Fig. 6a) are local image feature detectors, resembling the local fiber network structures. The filters in the other blocks (Fig. 6b-d) are more abstract, essentially representing various combinations of the local structures at different length scales and locations.

**Figure 6.**
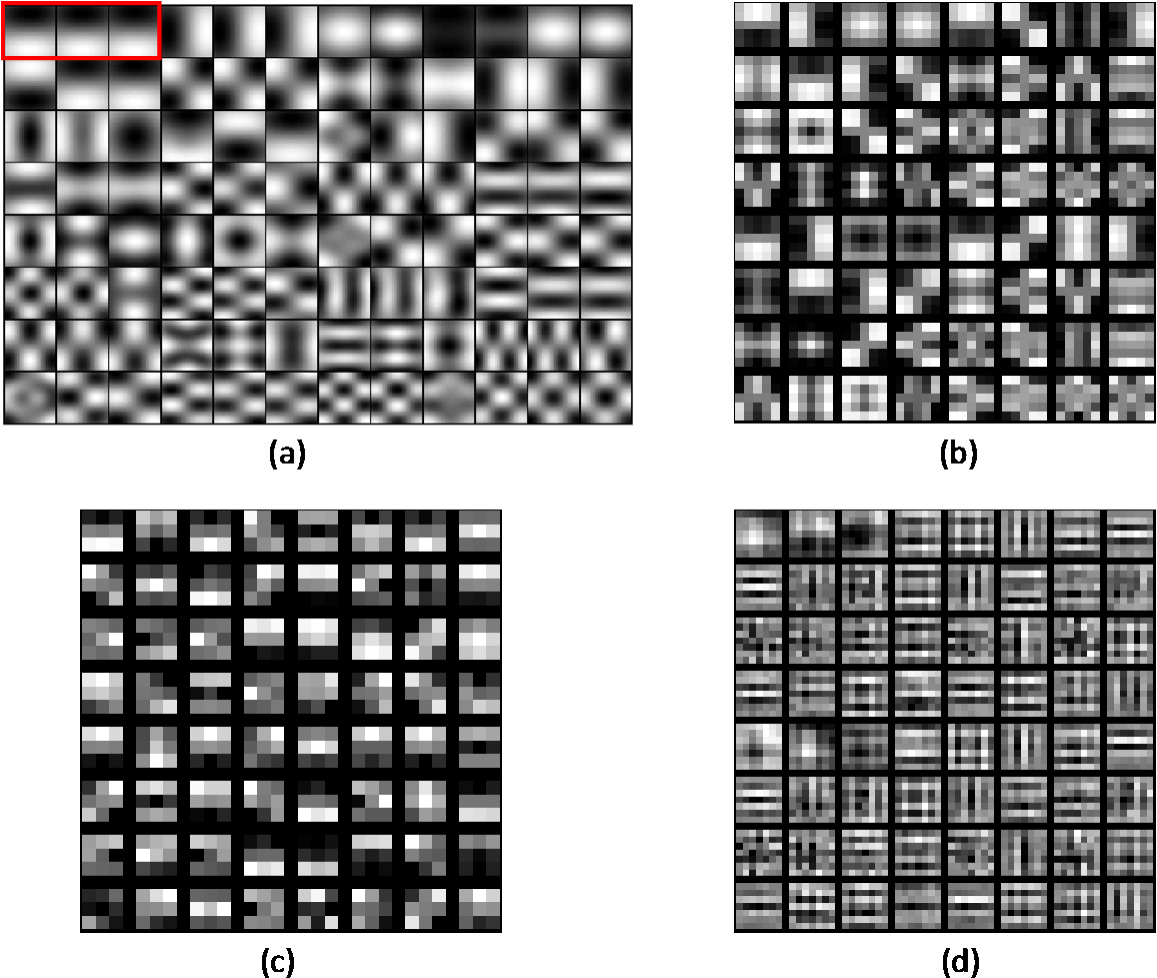
Examples of the learned filters. (a) The 32 filters in the convolution layer of the 2^nd^ block, the 32 opposites of these filters are not shown. The red box contains one filter (size is 33×33×3). (b) One of the filters in the convolution layer of the 3^rd^ block. (c) One of the filters in the convolution layer of the 4^th^ block. (d) One of the filters in the convolution layer of the 5^th^ block.

### 3.2 Classification

Classification performance was evaluated through ten-fold cross validation using the image patch data. In each round of cross validation, 90% of the image patches and corresponding overall stiffness values (i.e. sign of *α*_1_) were randomly selected as the training data; and the remaining 10% of the data were used as the testing data to test whether the trained classifier can predict the sign of *α*_1_, i.e., identify whether the tissue sample (corresponding to an image patch) is soft or stiff. The classification accuracy, defined as (TP+TN)/(TP+TN+FP+FN), the sensitivity, defined as TP/(TP+FN), and the specificity defined as TN/(TN+FP), were calculated to assess performance. Here, true positive (TP) is the number of stiff tissue patches correctly identified as stiff; false negative (FN) is the number of stiff tissue patches incorrectly identified as soft; true negative (TN) is the number of soft tissue patches correctly identified as soft; and false positive (FP) is the number of soft tissue patches incorrectly identified as stiff. In addition, AUC, defined as the area under a receiver operating characteristic (ROC) curve, was calculated as a measure of the overall classification performance. For comparison, a baseline softmax classifier using the skewness of image histogram [41] as the only feature, was also trained and tested. Since the two histograms of an image and its flipped version are the same, the flipped image patches were not used in the classification experiment. Two simplified versions of the CNN, CNN-s1 and CNN-s2 with less filters and less layers (details in Method section), were also tested.

ROC curves, as shown in Figure 7, were obtained by varying the discrimination threshold for each classifier. The performances of the proposed CNN, CNN-s1, CNN-s2, and the skewness-based softmax classifier using 0.5 as the discrimination threshold for classification, are reported in Table-1. The proposed CNN achieved the best performance, the skewness-based softmax classifier had the worst performance, and the two simplified CNNs had moderate performance.

**Figure 7.**
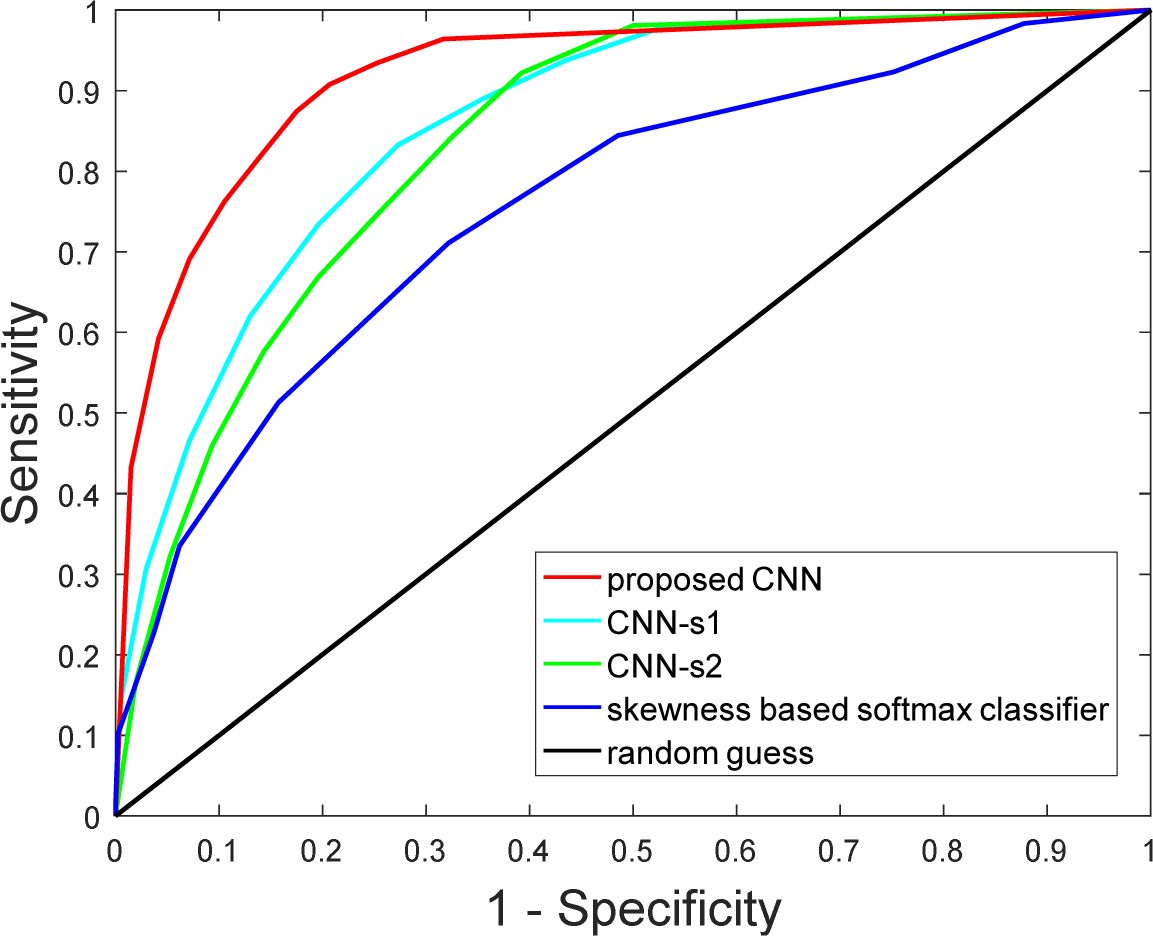
ROC curves of different classifiers

**Table-1:** Classification Performance

| Method | Accuracy | Sensitivity | Specificity | AUC |
| --- | --- | --- | --- | --- |
| proposed CNN | 84±2.5% | 82±4.1% | 86±3.6% | 0.92 |
| CNN-s1 | 78±2.8% | 73±5.5% | 80±3.7% | 0.86 |
| CNN-s2 | 75±3.5% | 67±5.6% | 80±4.9% | 0.84 |
| skewness based softmax classifier | 71±3.2% | 51±5.9% | 84±3.9% | 0.76 |

### 3.3 Regression

Regression performance was evaluated using a leave-one-out cross validation approach to test whether the trained regressor can predict the values of {*α*_1_, *α*_2_, *α*_3_}, which were used to reconstruct the stress-strain curve of each tissue sample by Eq.(1). In each round of the cross validation, the image patches and the stress-strain curves from one of the 48 tissue samples were used as the testing data to evaluate the accuracy of the regressor, and the remaining data were used as the training data to determine the parameters of the regressor. The predicted {*α*_1_, *α*_2_, *α*_3_} values for each of the image patches from the test tissue sample were averaged to obtain the final {*α*_1_, *α*_2_, *α*_3_} predictions for the whole tissue sample.

From the cross validation, the errors (Eq.(2)) in the predicted stress-strain curves were 0.021±0.015 and 0.031±0.029, compared to the actual S_11_~E_11_ and S_22_~E_22_ curves, respectively. Figure 8a shows an exemplary set of experimentally measured and predicted curves for one sample, and the error distribution across all of the samples is given in Figure 8b. The full set of predicted curves for all 48 samples are provided in the appendix.

**Figure 8.**
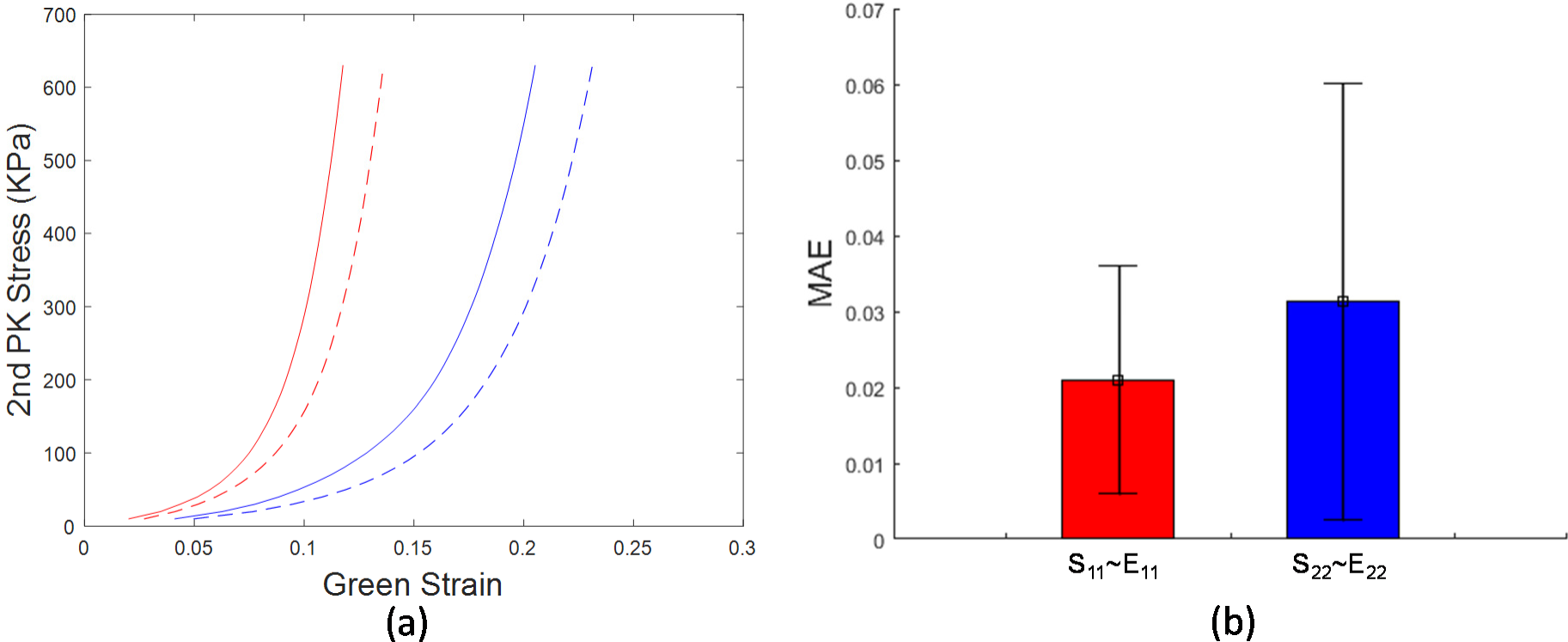
(a) Representative stress-strain curves predicted by the deep learning model shown as dashed lines, and the stress-strain curves obtained from mechanical testing shown as solid lines. S_11_~E_11_ curves are shown in red. S_22_~E_22_ curves are shown in blue. (b) The mean absolute error (MAE) distribution of all samples.

## 4. DISCUSSION

In this study, we developed a Deep Learning approach utilizing a deep CNN to estimate the elastic properties of collagenous tissues directly from noninvasive microscopy images. To our best knowledge, this is the first study in which Deep Learning techniques were used to derive nonlinear anisotropic elastic properties directly from tissue microscopy images. This work was motivated by the lengthy, complex, and destructive nature of traditional tissue mechanical testing. While it takes only about 10-30 minutes to obtain SHG images of a tissue sample, it takes much longer (hours) to prepare testing samples, set up testing and measurement instruments, and perform the actual mechanical test on each sample to obtain the stress-strain response curves. It took several months to obtain the data from the 48 samples used in this study. The success of this study holds promise for the use of Machine Learning techniques to noninvasively and efficiently estimate the mechanical properties of many structure-based biological materials.

Traditional machine learning methods [17] require hand-engineered features (i.e. features defined by human experts), which are difficult to obtain for this application. Rezakhaniha *et al.* [44] have defined intuitive texture features of tissue fibers, such as waviness, straightness, bundle size, etc., but this requires time-consuming manual annotation. Moreover, it is unclear whether these hand-engineered features could fully describe the fiber network structural information. As an end-to-end solution, CNN eliminates the need for hand-engineered features. One factor limiting the use of CNN and Deep Learning methods in biomechanics applications, is that they generally require a large amount of training data [42, 43], while the sample size for mechanical testing of biological tissues is typically small, on the order of 10 – 100 samples. However, in this study, it is shown that the deep CNN can also work well with a small dataset by combining supervised and unsupervised learning methods, and utilizing data augmentation methods. As more images and mechanical testing data are collected, the performance of the CNN can be further improved.

The CNN architecture used in this study, was specifically designed for this application. The 1^st^ to 5^th^ block of the CNN serve as automatic feature extractors that convert the input image into a feature vector for classification and regression. The filters in the first convolution layer represent different local fiber network patterns, while the filters in the remaining convolution layers represent various combinations of these patterns at different locations and length scales. Two simplified versions of the CNN were tested, i.e., CNN-s1 and CNN-s2 with less filters and less layers. The results show that simplifications to the CNN led to a significant decrease of accuracy, which may be the result of signal loss during signal propagation due to the fewer filters, and disruption of the encoding mechanism due to the fewer layers, respectively.

The CNN also demonstrated superiority over a simple image-feature based method to estimate the overall stiffness of collagen-based materials. Raub *et al.* [41] showed that the skewness of an image histogram was correlated to the collagen concentration and the Young’s modulus of collagen gels. Therefore, a softmax classifier was built by using the skewness as the only input feature in this study. As demonstrated in the results (Figure 7), the CNN outperformed the softmax classifier by a large margin; and even the two simplified versions of the CNN performed better than the softmax classifier, which underscores the superiority of CNNs for automatically extracting fiber network features.

More importantly, we demonstrated that the CNN can predict the PCA parameters of the stress-strain curves, such that the entire anisotropic stress-strain response of GLBP tissues can be estimated. For a nonlinear elastic response, it is well known that the Young’s modulus or stiffness cannot fully describe the tissue mechanical behavior, since the tangential value changes at different stress/strain levels along the nonlinear stress-strain curve. Thus, the PCA parameters offer a much more comprehensive look at the tissue elastic properties. Interestingly, we found that for this application, the overall “shape” of a stress-strain curve can be described with a single parameter, *α*_1_ in Eq.(1). The novel PCA based approach to represent stress-strain curves developed in this study may facilitate more thorough analysis and comparison of tissue stress-strain responses over basic stiffness metrics.

This approach opens the door for the fast and noninvasive assessment of collagenous tissue elastic properties from microstructural images, enabling many potential applications such as serving as a quality control tool for the manufacturing of BHVs.

## 5. CONCLUSION

In conclusion, this study demonstrated the feasibility of using Deep Learning techniques for fast and noninvasive assessment of collagenous tissue elastic properties from microcopy images. The main contributions of this study include: 1) the use of PCA to parameterize equi-biaxial stress-strain curves and quantify the overall stiffness, 2) the custom deep convolutional neural network design to automatically extract structural patterns of collagenous tissues, and perform classification to identify overall stiffness, as well as regression to predict PCA-parameters of nonlinear anisotropic stress-strain curves, and 3) the unsupervised deep learning method combined with supervised learning and data augmentation to overcome the challenge of small datasets for Deep Learning in the field of biomechanics. The developed approach was evaluated through cross validation, where an average classification accuracy of 84% and average regression errors of 0.021 and 0.031 were achieved. This study clearly demonstrates the great potential for Machine Learning techniques to estimate tissue mechanical properties solely through the use of noninvasive microcopy images.

## ACKNOWLEDGEMENTS

Research for this project is funded in part by NIH grant R01 HL104080. Liang Liang is supported by an American Heart Association Post-doctoral fellowship 16POST30210003. The authors thank Fatiesa Sulejmani and Andres Caballero for assisting in the collection of biaxial testing data and SHG images used in this study, as well as Caitlin Martin for comments and suggestions.

## Appendix

Predicted stress-strain curves of the 48 tissue samples are shown from the best to the worst. Horizontal axis shows Green Strain. Vertical axis shows 2^nd^ PK stress (KPa). Dashed lines are predicted stress-strain curves, and solid lines are the curves from mechanical testing.

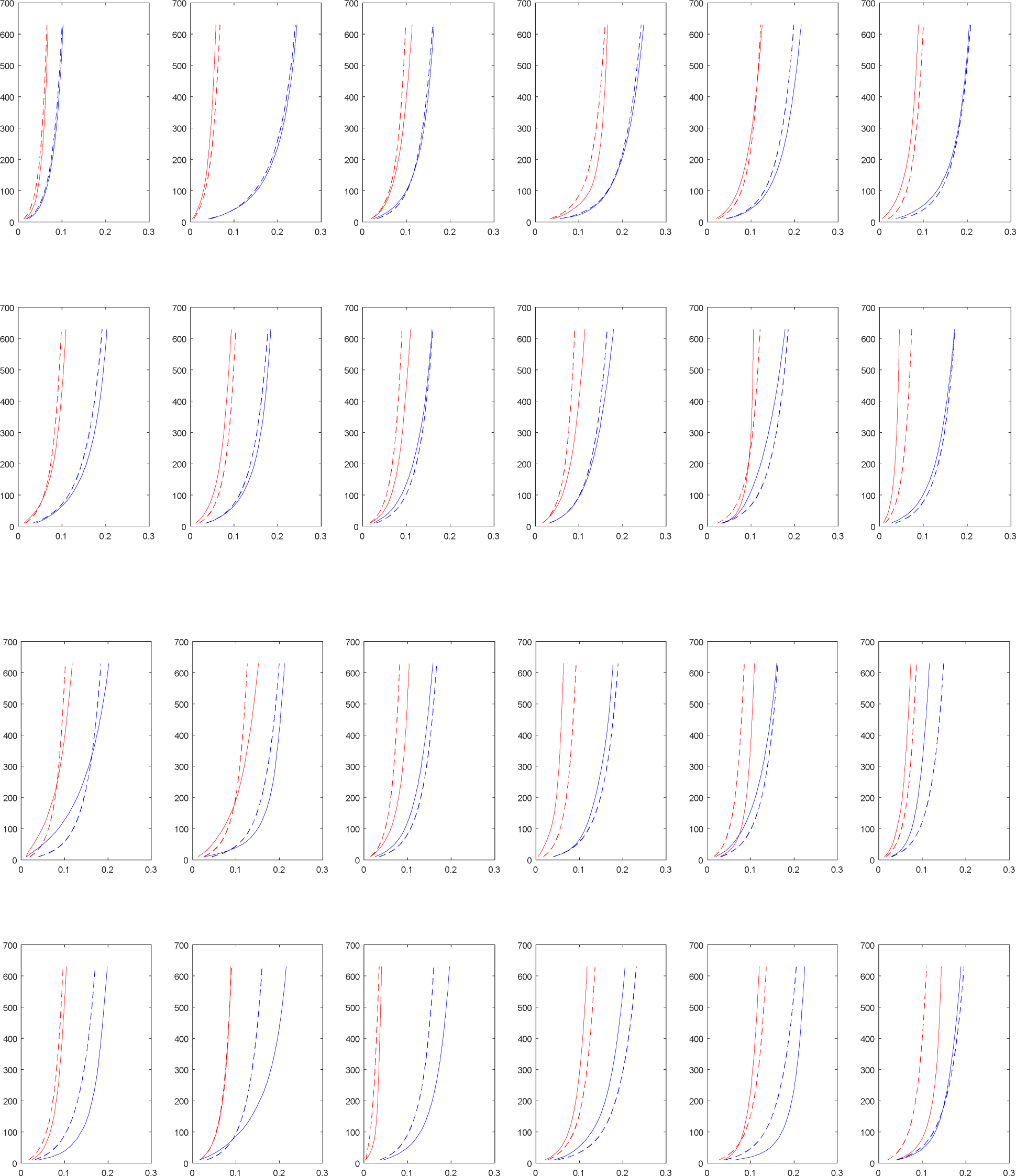

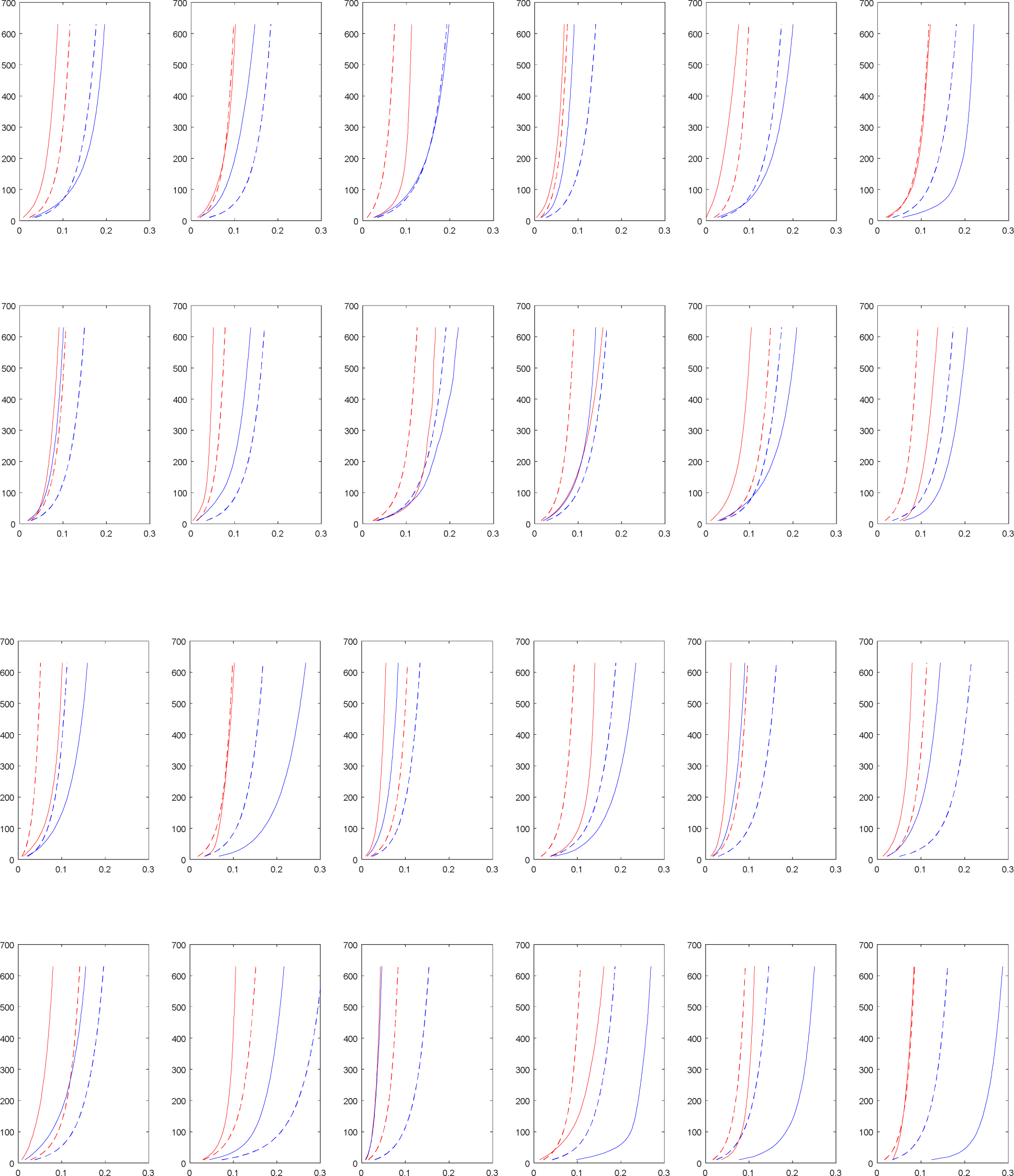

